# Oxidation and allocation of nectar amino acids during butterfly flight

**DOI:** 10.1101/2025.09.29.679119

**Authors:** Natasha Tigreros, Goggy Davidowitz, Chloe Bukholder, Chloe Chabaud

## Abstract

Flying animals face extreme energetic demands, relying mainly on carbohydrates and lipids, with occasional contributions from proteins and amino acids. In nectar-feeding species like butterflies and hummingbirds, sugars are the primary fuel, yet the extent to which other nectar-derived nutrients, like amino acids, are used for flight or retained for other functions remains unclear. Using ^13^C-labeled nectar, we tracked the metabolic fate of sugars and amino acids during flight in *Pieris rapae* butterflies. We found that proline and glycine, two of the most abundant nectar amino acids, were oxidized alongside sugars. Importantly, flight intensity modulated nutrient allocation from nectar: high-flight females incorporated less glycine into tissues, implying diversion toward flight, while threonine deposition in abdomens increased, reflecting prioritization for reproduction and storage. These findings reveal the complex role of nectar-derived nutrients in supporting locomotion and reproduction, while showing how nectar use can modulate trade-offs between flight and fecundity.

## INTRODUCTION

Powered flight is a key adaptation across animals, allowing insects, birds, and bats to forage, escape predators, locate mates, and disperse over long distances (Anderson & Ruxton, 2020; Dudley, 2002; Ruaux et al., 2020). Yet flight is also the most energetically costly mode of locomotion, far exceeding the metabolic demands of running or swimming (Bale et al., 2014; Schmidt-Nielsen, 1972). In insects, for example, flight can demand up to 100 times the resting metabolic rate (Anderson & Ruxton, 2020; Dudley, 2002; Ruaux et al., 2020). Thus, meeting these energetic demands requires efficient fuel use, with flying animals relying primarily on carbohydrates and lipids, and, in some cases, proteins and amino acids.The relative contribution of each fuel depends on the species, flight intensity, ecological context, and resource availability (Weber, 2011).

For nectar-feeding animals such as butterflies, bees, and hummingbirds, nectar sugars are the dominant energy source during flight (O’Brien, 1999; Suarez et al., 2011). Yet, nectar is a complex resource that also contains small but physiologically relevant amounts of amino acids (Baker & Baker, 1986; Nicolson & Thornburg, 2007). While studies, especially in insects, indicate that some amino acids are oxidized from internal stores during flight (Teulier et al., 2016; Stec et al., 2021; Weeda et al., 1979), it is still unclear whether amino acids obtained directly from nectar are similarly used in flight.

Beyond serving as a fuel for flight, nectar can also contribute to reproduction and long-term somatic functions. In many butterflies and moths, for example, nectar sugars as well as amino acids are incorporated into proteins and directly support oogenesis, supplementing or in some cases even replacing nutrient reserves acquired during larval development (Jervis & Boggs, 2005; Levin et al., 2017; Mevi‐Schütz & Erhardt, 2005). Similarly, in bird pollinators, especially those that rely heavily on nectar, amino acids in nectar have been suggested to help supplement their nitrogen requirements (Nicolson, 2007; Roguz et al., 2019). This dual role of nectar-derived resources may accentuate tradeoffs between flight and reproduction, as amino acids oxidized to sustain flight are no longer available for egg production. Thus, understanding whether nectar amino acids are metabolized during flight, or instead retained for tissue incorporation, is critical to link flight energetics with fitness and help clarify the physiological mechanisms underlying frequently observed trade-offs between flight and reproduction (Schmidt-Wellenburg et al., 2008; Tigreros & Davidowitz, 2019).

Here, we investigate the metabolic fate of nectar-derived nutrients in *Pieris rapae*, a common butterfly that relies on floral nectar throughout its adult life. Using ^13^C-labeled sugars and amino acids (proline, glycine, and threonine), we examined nutrient oxidation during flight as well as nutrient deposition in thoracic and abdominal tissues under varying flight conditions. By distinguishing which nectar components serve as immediate fuels and which are retained for other functions, we provide new insight into the role of nectar composition in supporting the energetics of insect flight.

## MATERIALS AND METHODS

### Study organism and Rearing

The cabbage white butterfly (*Pieris rapae*) is a widespread species whose adults feed on floral nectar, serving as pollinators for a variety of plants while obtaining sugars and amino acids as nutritional rewards (Alm et al., 1990; Corbera et al., 2018; Hongfang et al., 2010; Rader et al., 2009). Like other pollinators, female *P. rapae* can detect and preferentially consume nectars enriched in amino acids (Alm et al., 1990). This species faces high energetic demands due to its investment in both flight and reproduction: females fly an average of 0.7 km per day (Scott, 1987) and lay an average of 278 ± 172 eggs over a short adult lifespan of about 12.2 ± 4.0 days (Kimura & Tsubaki, 1986).

For this study, *P. rapae* were obtained from a laboratory colony established from a wild population in Utah, USA. Larvae were reared in a greenhouse on *Brassica oleracea*. Pupae and adults were maintained in a walk-in environmental chamber at 22 °C, 50% relative humidity, and a 16:8 h light–dark cycle.

### Nectar Treatments

In this study, we used artificial nectars that replicated the sugar and amino acid composition of *Lantana camara* (Verbenaceae), a high-quality nectar source frequently visited by diverse butterfly species. The composition of *L. camara* nectar has been well characterized and is widely used in studies of nectar preference in *P. rapae* (Alm et al., 1990) and other Lepidoptera (Erhardt & Rusterholz, 1998; Jervis & Boggs, 2005; Mevi-Schütz & Erhardt, 2003). Natural L. camara nectar typically contains ∼1 M sugars and ∼10 mM amino acids, yielding an approximate 1:100 amino acid-to-sugar ratio. Of the amino acid pool, roughly 6% comprises three essential amino acids, with threonine being the most abundant, while the remaining 94% consists of eight nonessential amino acids, dominated by proline and glycine.

To track use of specific nectar’s nutrients by female *P. rapae*, we created five lantana-mimicking nectar treatments. Four contained a single ^13^C-labeled compound—^13^C_1_-glycine, ^13^C_1_-proline, ^13^C_1_-threonine, or ^13^C-sucrose (enriched with cane sugar)—while the fifth served as a control, containing beet sugar and only unlabeled amino acids (δ^13^C ≈ –26.5‰). The labeled amino acids were selected because they are abundant in *L. camara* nectar and in other butterfly-pollinated flowers (Mevi-Schütz & Erhardt, 2003; Roguz et al., 2019). All isotope tracers were obtained from Cambridge Isotope Laboratories, Inc. (Tewksbury, MA, USA).

### Experiment 1. Oxidation of nectar nutrients during flight

To assess oxidation of nectar nutrients during flight, females were fed 48 h post-emergence with 10 µl of either a Control or one of four ^13^C_1_-labelled nectars: ^13^C_1_-glycine, ^13^C_1_-proline, ^13^C_1_-threonine, or ^13^C-sucrose. Four hours after feeding, each female was placed in a respirometry chamber consisting of a 500 ml syringe fitted with an injection port and adjusted to provide a 200 ml chamber volume. This volume was sufficient to allow flight while enabling CO_2_ accumulation to ≥300 ppm. The syringe air was replaced with dried, CO_2_-free air, sealed, and females were flown for 4 min under natural light, with gentle shaking to prevent resting. A 20 ml breath sample was then analyzed for δ^13^CO_2_ using a cavity ring-down spectrometer (G2201-i, Picarro Inc., Sunnyvale, CA) coupled to a SSIM2 Small Sample Isotope Module (Picarro Inc., Sunnyvale, CA).

### Experiment 2. The effect of flight on nectar nutrients depositions

To determine where nectar-derived amino acids are deposited, we used a different group of adult females (48 h post-emergence) which were randomly assigned to one of two flight treatments— Low or High flight intensity—and fed either a control nectar or one of four ^13^C_1_-labelled nectars. For the Low flight intensity group, females were kept in a 30 × 30 × 30 cm mesh cage lined with paper towel inside a climate-controlled walk-in chamber (LD 16:8 h, 22°C at 50% RH). For the High flight intensity group, females were kept under the same conditions but were additionally stimulated to fly once per day, for five continuous minutes, by gently touching them with a fine paintbrush (Niitepõld & Boggs, 2015). Preliminary trials showed that longer flight durations caused extreme exhaustion, with butterflies becoming unresponsive. All females were fed once daily with 10 µl of their assigned nectar treatment. After three consecutive days of the flight and nectar treatments, individuals were euthanized by freezing at −20 °C. Thoraces and abdomens were dissected, dried at 50 °C for 48 h, and homogenized. δ^13^C values were then measured using cavity ring-down spectroscopy (CRDS; Picarro stable isotope analyzer, G2201-i) coupled with an A0201 combustion module and A0301 gas interface.

### Statistical Analysis

To assess nectar-derived nutrient oxidation (Experiment 1) and tissue deposition (Experiment 2), we used Dunnett’s test to compare the δ^13^C values of breath or tissue samples from females fed each ^13^C-labeled nectars against those of control females fed unlabeled nectar.

Because δ^13^C values can naturally change with tissue type and treatment, we used δ^13^C of control females to estimate atom % excess (APE), which is a unit of enrichment (McCue et al., 2011; Slater et al., 2001). We used ^13^C concentrations of both control and labelled tissues expressed in δ^13^CVPDB to calculate the atom percent (AP) and then Atom Percent Excess (APE), defined as *APE = AP(label) – AP(control)* (Slater et al., 2001).

The effects of flight on nutrient allocation to thoracic and abdominal tissues were analyzed using two-way ANOVAs with flight treatment, nectar nutrient treatment (^13^C-labeled vs. control), and their interaction as fixed factors. When significant interactions were detected, we estimated predicted values and performed pairwise comparisons of simple effects using the *emmeans* package in R, applying a Tukey adjustment for multiple testing. All statistical analyses were conducted in R 3.6.0. and assumptions of normality and homogeneity of variance were verified prior to analysis.

## RESULTS AND DISCUSSION

Nectar is a critical resource for flying pollinators, providing the energy needed to support their high metabolic demands during flight (Brien, 1999; Suarez, 2005; Welch et al., 2006). While it is well established that nectar sugars serve as the primary fuel for flight, far less is known about the roles of non-sugar components, like amino acids. Here we investigated the use of nectar-derived nutrients, including essential and non-essential amino acids, as metabolic fuels during flight and how flight activity may influences their allocation to different tissues in female *Pieris rapae*.

### Oxidation of nectar nutrients during flight

Females that fed on nectar containing ^13^C-labeled sugars, proline, or glycine showed significant ^13^CO_2_ enrichment (more positive δ^13^C values) in breath compared to controls fed unlabeled nectar (**Fig. 1**), indicating active oxidation of both sugars and non-essential amino acids. Although previous studies have suggested that nectar amino acids help meet the high metabolic demands of flying nectarivores (Carter et al., 2006; Levin et al., 2017; Stec et al., 2021), our results provide the first direct evidence that nectar amino acids are actually oxidized during flight. At the same time, no significant oxidation of threonine by flying females was observed **(Fig. 1)**, suggesting that essential amino acids may be less likely to contribute to locomotion, at least under our experimental conditions. These results add to the growing body of evidence highlighting the importance of nectar amino acids for flying animals, and in particular for insect pollinators. Proline and glycine are common constituents of floral nectar (Baker & Baker, 1973; Rusterholz & Erhardt, 1998) and can act as a phagostimulant in pollinator insects species (Lim et al., 2019; Ruedenauer et al., 2019). Several studies have shown that proline, in particular, is selectively oxidized in insect flight muscle, especially during the initial 30 seconds of flight (Stec et al., 2021; Teulier et al., 2016; Weeda et al., 1979). Though glycine’s role in flight is less defined, its consumption has been linked to neuromodulation and memory in insects (Parkinson et al., 2025), suggesting a dual function: fuelling flight and supporting cognitive processes essential for effective foraging.

**Fig. 1.**
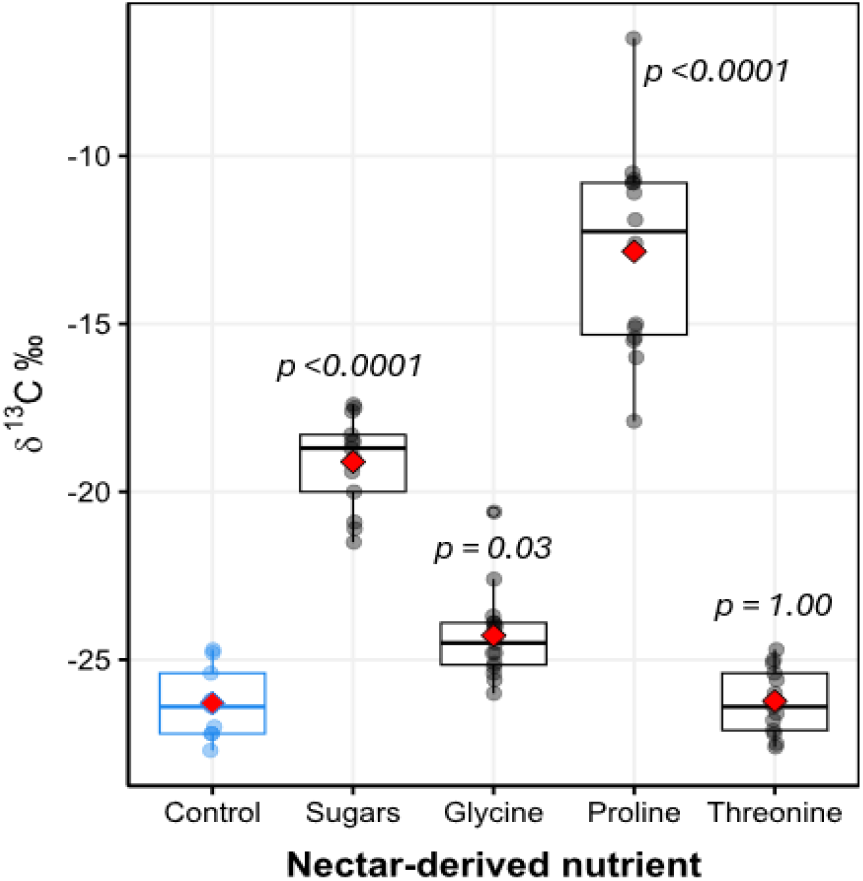
Boxplots show the *δ*^13^C values from exhaled breath during flight of *P. rapae* females. Significant differences between the δ1^3^C from control (n = 9, blue boxplot) and females that consumed ^13^C-labelled nectars indicate that a nectar-derived nutrient was oxidized during flight. Traced nutrients included sugars (n = 13), two non-essential amino acids: glycine (n = 15) and proline (n = 14), and an essential amino acid, threonine (n = 13).

### The effect of flight on nectar nutrients depositions

Gravimetric measurements of the impact of flight on thorax and abdomen resources did not indicate any differences (both thorax and abdomen *p >* 0.05). In contrast, isotopic analysis of control, unlabeled females, revealed significant changes in tissue carbon composition due to flight intensity. Although abdominal tissue had higher δ^13^C values than thoracic tissue (ANOVA: F_1,20_ = 8.1, P = 0.01), females subjected to high-intensity flight exhibited overall ^13^C enrichment in both thorax and abdomen compared to low-flight individuals (F_1,20_ = 6.26, P = 0.02), consistent with increased oxidation and depletion of lipid stores (**Fig. 2**).This is in line with previous studies showing that butterflies rely on both carbohydrates and lipids to meet the high energy costs of flight, and that sustained flight can rapidly deplete internal energy, often leading to flight fecundity tradeoffs (Tigreros & Davidowitz, 2019).

**Figure 2.**
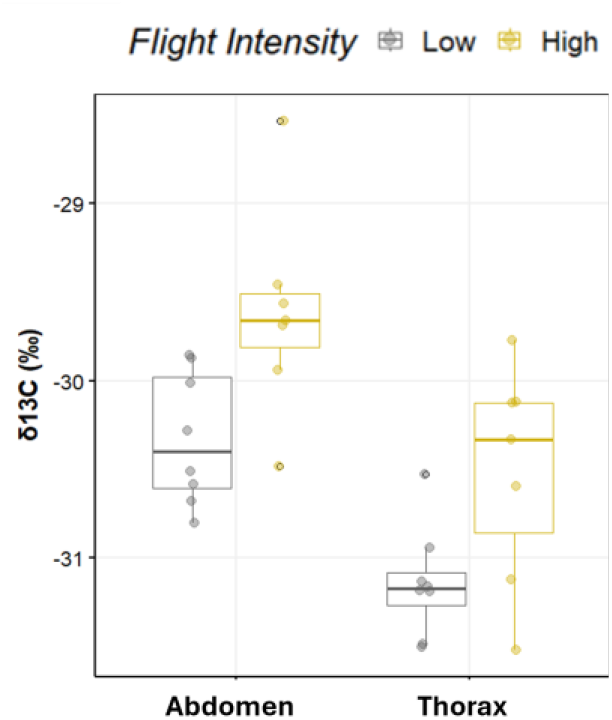
Changes in δ^13^C values in thorax and abdomen tissues of *Pieris rapae* females subjected to Low versus High flight intensity. Elevated δ^13^C values indicate ^13^C enrichment, reflecting relatively lower lipid content in abdomen tissue (compared to thorax) and highly active individuals.

Previous studies have shown that nectar in general and its amino acids in particular can be rapidly incorporated, within minutes, into Lepidoptera flight muscle and developing oocytes (DeFino & Davidowitz, 2024; Levin et al., 2017). Given that the thorax serves as the primary site of fuel combustion during flight (Dudley, 2002; Treidel et al., 2024), it can be expected that incoming amino acids and sugars are preferentially oxidized rather than retained. Here, we demonstrate that flight activity significantly alters the allocation of nectar nutrients across female tissues (*F*_3, 102_ = 5.33, *P* = 0.002). In the thorax, the presence of nectar-derived nutrients declined with increasing flight intensity (*F*_1,51_ = 4.99, *P* = 0.03; **Fig. 3**), consistent with their increased metabolic oxidation or mobilization. Interestingly, while most labelled amino acids were detected in both thorax and abdomen tissues (**Fig. S1**), proline was notably absent in the thorax under high flight conditions, and it only approached significance in low-flight males (*P* = 0.09; **Fig. S1**). This supports the hypothesis that proline is rapidly oxidized during flight, leaving little available for tissue incorporation.

**Fig. 3.**
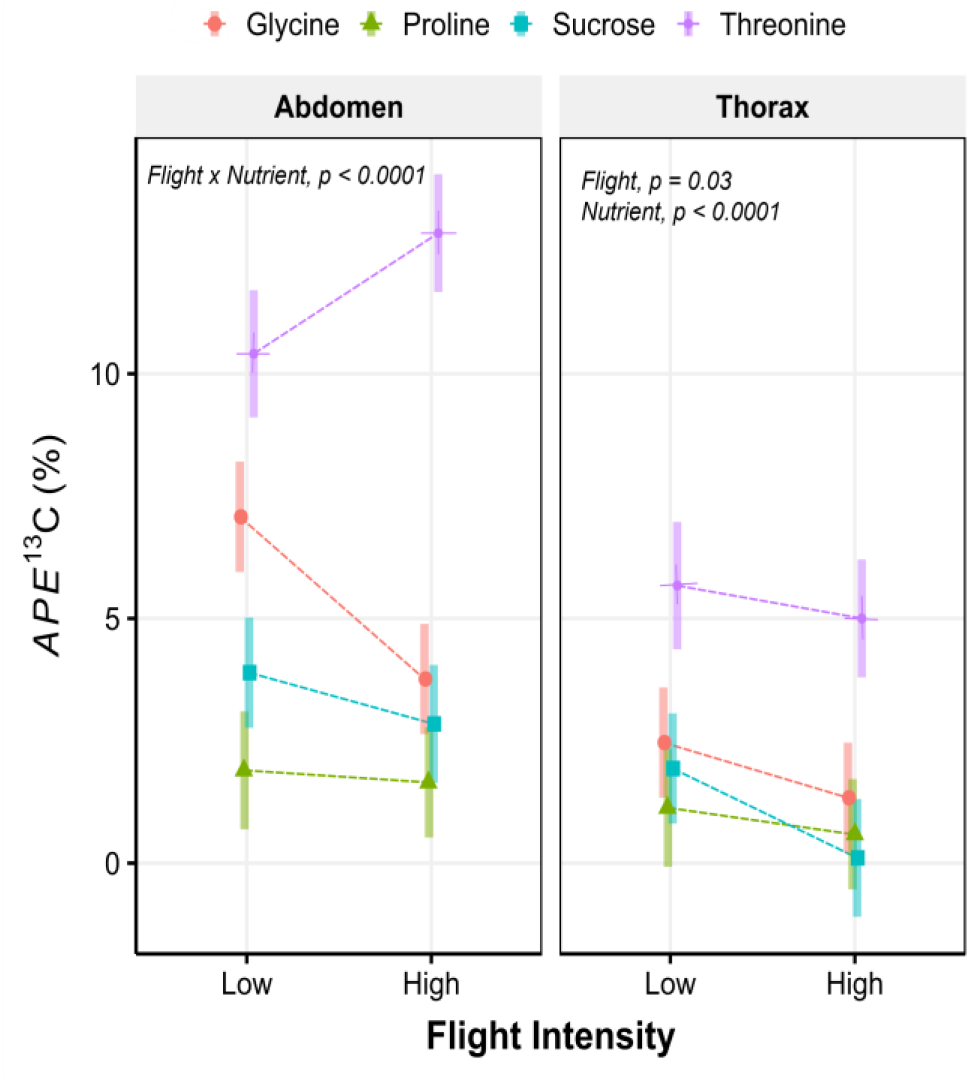
Effects of flight, high vs. low intensity, on the use of nectar-derived nutrients in abdomen (left panel) and thorax (right panel) tissues of *P. rapae* females. The y-axis represents the estimated marginal means ± 95CI of the atom percent excess (APE ^13^C (%)), with higher APE values indicating greater incorporation of nectar derived sucrose, non-essential amino acids (glycine and proline) and an essential amino acid (threonine).

Patterns of nutrient incorporation and use were distinct in the abdomen, reflecting its key roles in nutrient storage and reproduction (**Fig. 3**). We found that the incorporation of nectar nutrients varied by type (*F*_3,51_ = 4.9, *P* = 0.01): glycine decreased (p = 0.002), while threonine increased (p = 0.03) with higher flight activity **(Fig. 3; Table S1)**. Interestingly, proline and sucrose incorporation remained unchanged (**Fig. 3; Table S1**). As the primary site of nutrient storage and reproduction, the abdomen likely channels nectar-derived nutrients toward oogenesis and long-term metabolic demands (Boggs, 2009; O’Brien et al., 2002). The decline in abdominal glycine, an amino acid significantly oxidized during flight (**Fig. 1**), suggests that this non-essential amino acid may be diverted to meet immediate energy demands, possibly contributing to a trade-off with female fecundity. Conversely, the increase in threonine, an essential amino acid, may reflect its selective prioritization for egg provisioning. These findings support the idea that adult females use nectar to buffer against nutrient limitations, whether from poor larval diets (Mevi-Schütz & Erhardt, 2005; O’Brien et al., 2000) or increased metabolic demands from flight.

Collectively, our findings demonstrate that a flying nectarivorous insect utilizes both sugars and specific amino acids—particularly proline and glycine—as metabolic fuels during flight. Moreover, intense flight alters the fate and allocation of nectar-derived nutrients in adult tissues, likely reflecting a trade-off between immediate energy needs and resource storage. Although direct evidence in nectarivorous vertebrates is scarce, the high concentrations of amino acids (including proline and glycine) in nectar of some bird-pollinated plants suggest these compounds may serve similar functions in these species (Nicolson, 2007; Roguz et al., 2019). Thus, our results provide broader insights into how nectar composition can influence locomotion and fitness of nectar-feeding species, emphasizing the ecological significance of amino acids in adult diets, especially under conditions of elevated energy expenditure.

