## Supplementary material for "Oxidation and allocation of nectar amino acids during butterfly flight": Table S1 and Figure S1

**Table S1** (A) Statistical output from general linear models (Anova type III) on the effects of flight intensity (high vs low) on deposition of nectar-derived nutrients (Nutrient) abdomen and thorax of *P. rapae* female. A significant interaction between flight and nutrient was further investigated by (B) estimating predicted values and conducting pairwise comparisons of their simple effects. Degrees of freedom are indicated in parenthesis [d.f.]

(A) Anova Summary Table

| | **Abdomen** | | **Thorax** | |
|---|---|---|---|---|
| | *F* | *p* | *F* | *p* |
| Flight [d.f.= 1, 51] | 10.60 | < 0.01 | 4.99 | 0.03 |
| Nutrient [d.f.= 3, 51] | 22.16 | < 0.0001 | 24.65 | < 0.0001 |
| Flight x Nutrient [d.f.= 3, 51] | 4.90 | < 0.01 | 1.21 | 0.31 |

(B) Simple effects: **Abdomen**

| | High vs. Low Flight | |
|---|---|---|
| | *estimate* | *p-value* |
| Sucrose | 1.05 | 0.32 |
| Glycine | 3.33 | 0.002 |
| Proline | 0.25 | 0.81 |
| Threonine | -0.246 | 0.03 |

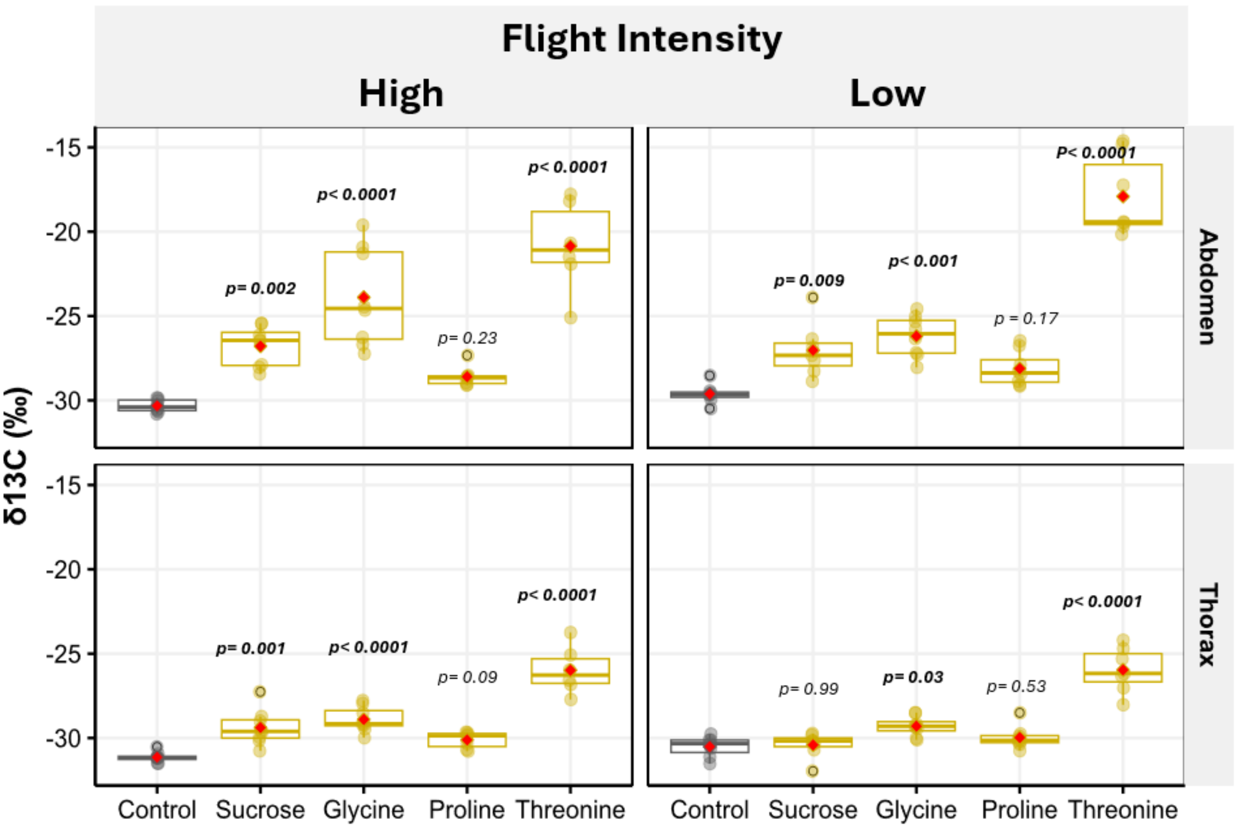


**Figure S1** Boxplots with arithmetic mean (red diamon) show the $\delta^{13}$C values from the thorax and abdomen tissues of female *P. rapae* that consumed $^{13}$C-labelled nectar compared to controls that consumed unlabeled nectar. The figure is divided into four panels based on body part and flight intensity level (high vs. low). The $^{13}$C -labeled nectar treatments were composed of one of four nutrients commonly found in *Lantana camara* nectar and other butterfly pollinated plants. These included two non-essential amino acids (glycine and proline), one essential amino acid (threonine), and sucrose. p-values denote significant differences, from Dunnett test, between females fed a $^{13}$C-labelled nutrient with its respective control (grey boxplot).
